# Why the tumor cell metabolism is not that abnormal

**DOI:** 10.1101/865048

**Authors:** Pierre Jacquet, Angélique Stéphanou

**Author notes:** Corresponding author’s.

## Abstract

The cell energy metabolism is a multifactorial and evolving process that we address with a theoretical approach in order to decipher the functioning of the core system of the glycolysis-OXPHOS relationship. The model is based on some key experimental observations and well established facts. It emphasizes the role of lactate as a substrate, as well as the central role of pyruvate in the regulation of the metabolism. The simulations show how imposed environmental constraints and imposed energy requirements push the cell to adapt its metabolism to sustain its needs. The results highlight the cooperativeness of the two metabolic modes and allows to revisit the notions of *metabolic switch* and *metabolic reprogramming*. Our results thus tend to show that the Warburg effect is not an inherent characteristic of the tumor cell, but a spontaneous and transitory adaptation mechanism to a disturbed environment. This means that the tumor cell metabolism is not fundamentally different from that of a normal cell. This has implications on the way therapies are being considered. The quest to normalize the tumor acidity could be a good strategy.

**Author Summary:** Cancer cells metabolism focuses the interest of the cancer research community. Although this process is intensely studied experimentally, there exists very few theoretical models that tackle this issue. One main reason is the extraordinary complexity of the metabolism that involves many inter-related regulation networks which makes it illusory to recreate computationally this complexity. In this study we propose a simplified model of the metabolism which focuses on the interrelation of the three main energetic metabolites that are oxygen, glucose and lactate with the aim to better understand the dynamic of the core system of the glycolysis-OXPHOS relationship. However simple, the model highlights the main rules that allow the cell to dynamically adapt its metabolism to its changing environment. It moreover allows to address this impact at the tissue scale. Simulations performed in a spheroid exhibit non-trivial spatial heterogeneity of the energy metabolism. It further reveals that the metabolic features that are commonly assigned to cancer cells are not necessarily due to cell intrinsic abnormality. They can emerge spontaneously because of the disregulated over-acidic environment.

## Introduction

The energy metabolism of cancer cells has been the subject of extensive research for over fifty years, yet the mechanisms governing tumors metabolism are not clearly understood. The Warburg effect, which now seems accepted as a key feature of many types of cancer, is considered by some as one possible fundamental cause of cancer [1, 2]. Some define this effect as a high lactate production despite sufficient oxygen supply [1, 3]. However according to Warburg’s original observations in the 1920s, the effect is limited to the production by the tumor of a large amount of lactate (independently of the oxygen presence) [4–6]. This lactate production is induced by a high glycolytic activity and increased glucose uptake. This creates around the cells, and especially within solid tumors, a whole microenvironment, characterized by an acidic pH, favoring tumor cells invasion. One recurring question remains “how do these extreme conditions benefit the cell ?” [7–9]. Understanding the impact of the microenvironment on ATP production may be part of the answer.

The purpose of this paper is to tackle this issue with a theoretical approach to bring some new understanding of the cell energy metabolism.

The cell metabolism is highly complex since it is a multifactorial mechanisms that involves many different interacting processes with many different actors. Moreover it is an evolving process and although crucial, this aspect is rarely considered and often overlooked. In this context, a theoretical model is a powerful and efficient way to make sense of this complexity and to address the temporality. It allows to test the pertinence of some new hypotheses and to exhibit some emergent properties that cannot be intuited, so as to provide a better understanding of the intimate functioning of the metabolic machinery and also to provide new insights to guide future research.

Several models have been proposed to describe the cell energy metabolism [10–14] but some can be too complex to be easily reused and tested by experimentation. We therefore focused more specifically on models that describe the production of ATP according to the conditions surrounding the cell. Extracellular oxygen and glucose concentrations, lactate production and quantification of the extracellular pH (by protons secretion) are the conditions that have been mainly considered in modelling. The availability of glucose and oxygen respectively influences the activity of glycolysis and oxydative phosphorylation (OXPHOS). Casciari et al. [15] proposed a model that describes glucose and oxygen consumption changes in EMT6/Ro cells. They raised the importance of pH on these uptakes and mathematically formalized these observations. This pioneering model − that exploits experimental data − has since been used in many studies [11, 16, 17].

Our computational model is again primarily inspired by this reference model. However, it additionally integrates the most recent knowledge, in particular about the disappearance of the Warburg phenotype under acidic conditions [18], and is rooted on some new key observations and established facts. The model focuses once more on the glycolysis-OXPHOS relationship but emphasizes the role of lactate as a substrate. Lactate is indeed of particular interest since it can allow the tumor cells to survive despite a significant depletion of glucose [19]. Its role for cell viability under acidic conditions have been overlooked since very few models integrate this important fact.

Our model also takes into account the central role of pyruvate in the regulation of the metabolism. It might seem trivial to recall that, glycolysis is the set of reactions that allows to transform glucose into pyruvate, but sometimes glycolysis is mentioned as the complete transformation of glucose to lactate (also called fermentation). This is a source of misunderstanding, since it suggests that glycolysis is opposed to the mitochondrial metabolic pathway that would be an independent process. In fact the Krebs cycle and the subsequent oxidative phosphorylation (OXPHOS) requires pyruvate from glycolysis in normal conditions, as a first step, hence its pivotal role. These paths, although running in parallel in the cell, are in reality a chain of reactions and not dual options that exclude one another.

The model then investigates how imposed environmental constraints and imposed cell energy requirements push the cell to adapt its metabolism to sustain its needs. The simulations performed are insightful since they clearly show how glycolysis and OXPHOS are used concomitantly and in a cooperative way [20]. The gradation in their relative contributions to ATP production is shown to depend on the available resources and environmental acidity. The results also allow to revisit the notions of *metabolic switch* and *metabolic reprogramming*.

The model of cell energy metabolism may appear over-oversimplified considering the huge complexity of the fine regulation mechanisms at work. We did not consider the role of glycolysis in the biosynthesis of amino-acid either. We want to stress the point that at this stage our goal was to focus on the global behavior of energy metabolism to be able to address its repercussions at the tissue scale and to highlight spatial metabolic heterogeneity in a spheroid.

## Results

### Integration of the latest experimental evidences for a new understanding of cell energetic metabolism: focus on the pivotal role of pyruvate

To understand the nature of the Warburg effect, it is necessary to define the energy mechanisms that govern the cell. One of the questions that may arise is whether the Warburg effect appears as an inherent tumor cell characteristic or, on the contrary, as a transitory behavior of the cell’s metabolism in response to the environmental constraints. In other words is it appropriate to continue to describe this phenomenon as a *metabolic switch* [21, 22]? When considering the metabolism of a cell, many parameters must be taken into account, as several metabolic pathways are involved. Here we focus on the glycolysis-OXPHOS system, as well as on the place of lactate within it. It is nevertheless important to remember that other pathways such as *β*-oxidation of fatty acids or glutaminolysis can contribute to increase the reaction intermediates and thus increase the energy production capacity.

In order to define our model, we think that it is useful to recall some fundamental concepts of the metabolism of these two pathways. Here glucose is considered as the main source of energy in the cell. This molecule is catabolized during a sequence of three essential processes in order to produce ATP. The first reaction, glycolysis, transforms glucose into pyruvate as follows:

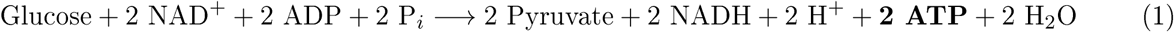

NAD^+^ is rate limiting for glycolysis, in the reaction catalysed by Glyceraldehyde 3-phosphate dehydrogenase (GAPDH). But in the overall process of fermentation / respiration, the NAD^+^ pool is refilled through LDH, or oxydative phosphorylation. If the ratio NAD^+^ / NADH is too low the glycolysis will be inhibited. Many metabolic reactions modify this ratio and more generally the redox state of the cell. Here the goal is not to model the set of mechanisms that can lead to change the ratio NAD^+^ / NADH. At the moment we therefore consider that NAD^+^ is not limiting for the processes we seek to observe, although it is important to be aware that this can have a significant impact on energy metabolism.

Pyruvate can be reduced to lactate with lactate dehydrogenase (LDH) but the reaction will not produce more ATP. Pyruvate can also be decarboxylated by the pyruvate dehydrogenase in Acetyl-CoA (Fig.1). This decarboxylation takes place in the mitochondria.

**Figure 1:**
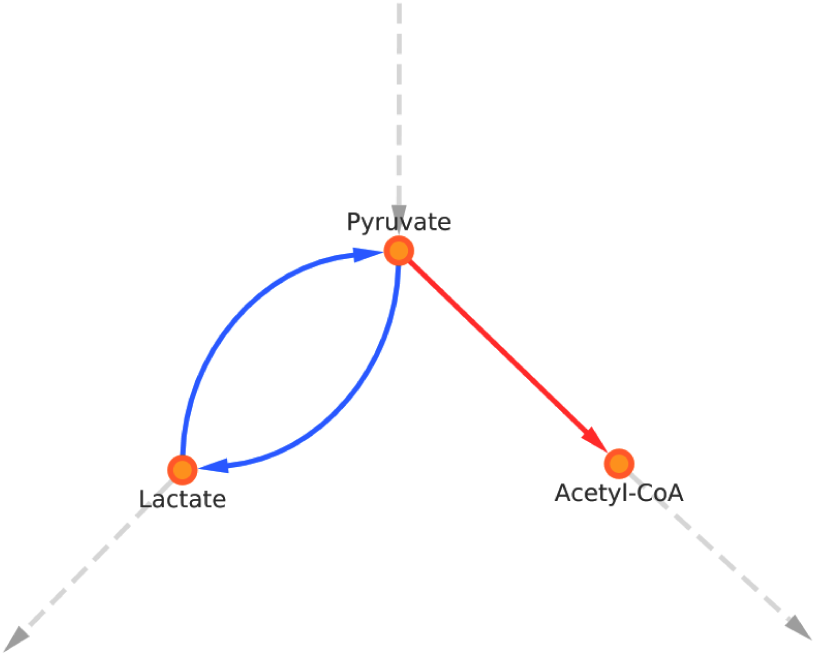
Fate of pyruvate. It can be reduced to lactate or decarboxylated in Acetyl-CoA. The pyruvate-lactate reaction is reversible, pyruvate can be restored from lactate.

If the pyruvate is converted into lactate, the latter, under physiological conditions, will be secreted in the extracellular space by MCT transporters. If not, the pyruvate is converted to Acetyl-CoA, that enters the citric acid cycle, generates one GTP (equivalent to one ATP) and generates NADH and FADH_2_ used in the final step of the reaction. It is the OXPHOS, in which the energy released by the transfer of electrons from a donor to an acceptor (oxygen in particular), that is used to produce a large quantity of ATP (from ADP). This final reaction can be summarized as follows:

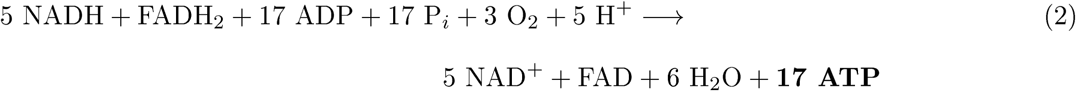

Each reactions are written in their canonical form. From one molecule of glucose, the cell can obtain a total of 38 ATP molecules. Currently, the estimate of the number of ATP molecules produced during aerobic respiration is still under debate (38 is a theoretical maximum) [23, 24]. We will continue our calculation with 17 moles of ATP per mole of pyruvate [16]. It is also relevant to note that in several experiments the level of pyruvate remains relatively constant (regardless of the microenvironment) [18]. We therefore hypothesize that pyruvate functions as a reservoir that flows according to the mitochondrial energy requirement. If this reservoir overflows (too much pyruvate produced), the excess is converted into lactate. Conversely, if it empties faster than it fills (not enough pyruvate), production or consumption can be readjusted (by reabsorbing lactate for example). We note that PKM2 (Pyruvate Kinase) enzyme is limiting in the final reaction of pyruvate production. This enzyme is tightly regulated and this regulation determines if glycolytic intermediates before pyruvate should be used in synthesis of amino-acid/nucleic acid or not. In a cancer scenario, the PKM2 enzyme is mainly in its inactive dimeric form but can switch to its trimeric form by the accumulation of Fructose 1,6-bisphosphate (FBP) which leads to convert most of glycolytic intermediates to pyruvate [25]. The tetramer/dimer ratio of PKM2 enzyme oscillates [26]. This mechanism is not studied here but might change the temporal dynamic of glycolysis. However, the fact that a lot of lactate is produced in cancer cells, indicates that on a longer time period, the PKM2 enzyme still allows the reaction to proceed. Pyruvate can also be produced from oxaloacetate by pyruvate carboxylase to remove an excess of oxaloacetate in the TCA cycle. Finally pyruvate is also used to produce alanine. These two mechanisms are not considered in this model.

The model is based on the following experimental observations:

1. glucose consumption increases with glucose concentration, oxygen consumption increases with oxygen concentration [15, 27] and lactate consumption increases with lactic acidosis [18, 28];
2. the less oxygen in the extracellular medium, the more glucose consumption increases [29] up to a saturation threshold [15];
3. the more acidic the medium (the protons concentration is high), the lower the glucose uptake [15, 18, 30, 31];
4. pyruvate concentration remains relatively constant thanks to pH variations [18].

From this experimentally established basis, several metabolic scenarios emerge. Three representative scenarios are presented in figure 2.

**Figure 2:**
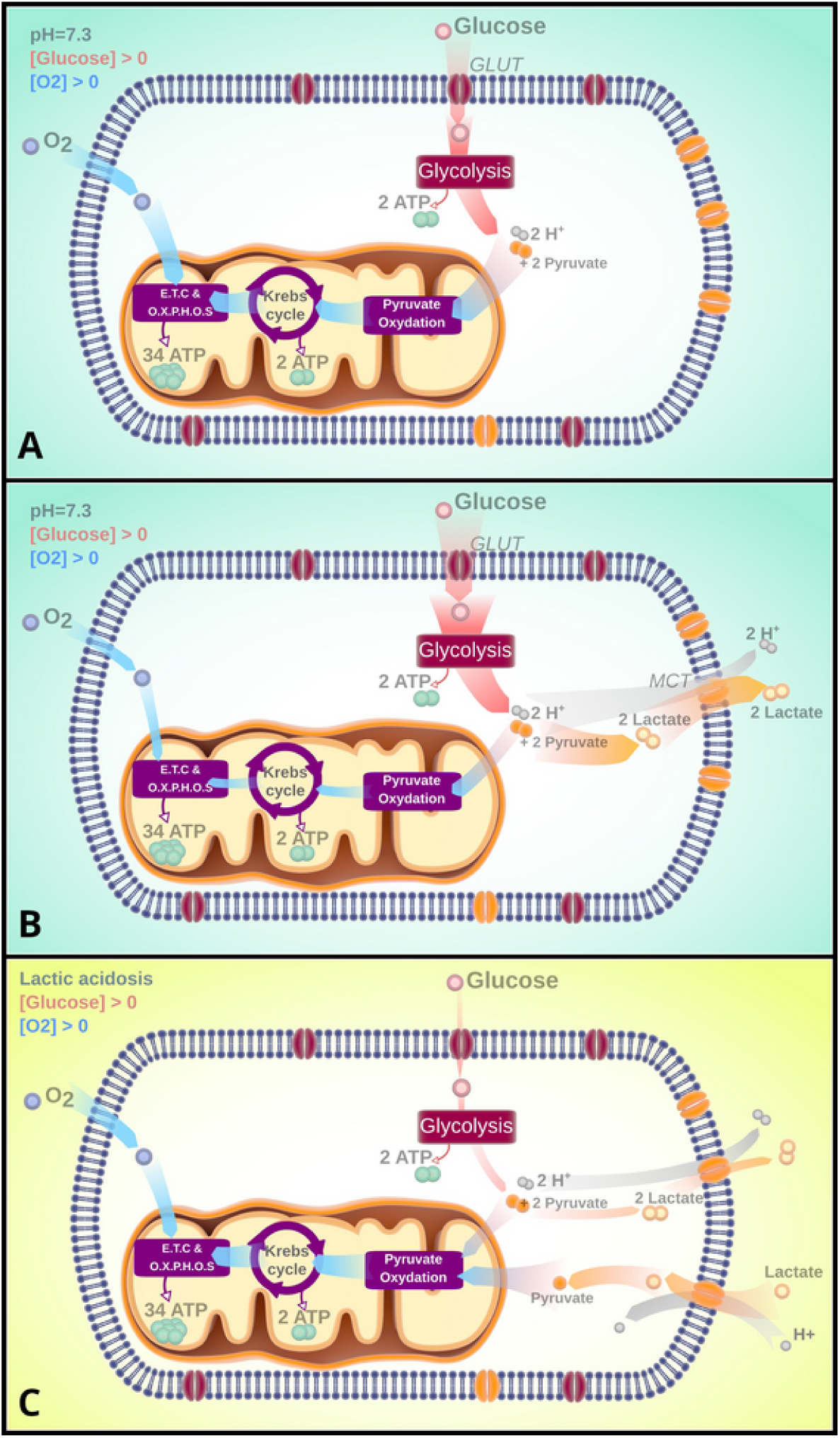
**A.** Healthy cell − The majority of the glycolytic flux is redirected in the mitochondria, and little or no pyruvate is converted to lactate; **B.** Warburg effect − The glycolytic flux is higher than the level that allows the balance between the production of pyruvate and its consumption. As a result, the pyruvate is partly converted into lactate and protons are Secreted and progressively acidify the medium; **C.** Lactic acidosis − When the pH is low the glycolytic flux drops. There is not enough pyruvate to supply the OXPHOS, so the net flux of lactate enters the cell and is converted back into pyruvate.

Figure 2A summarizes the different reactions expressed in (1) and (2), in a healthy cell under physiological conditions. Figure 2B, shows a cell with a Warburg phenotype. As in figure 2A, this cell uses both glycolysis and OXPHOS, but the glycolytic flux is higher (or the ATP demand is lower), reducing the need to produce ATP by the mitochondria. Pyruvate accumulates, shifting the flux to lactate production. These mechanisms push the cell towards the scenario presented in figure 2C. In this case, as protons and lactate accumulate, the glycolytic flux is substantially reduced [18, 28], increasing the mitochondrial activity and increasing the demand for pyruvate. The net flow of lactate enters the cell and is converted back into pyruvate to maintain the cellular level. Here the Warburg effect stage, is not the final stage of the tumor cell, but a transient one.

To estimate the rate of ATP production per cell, it is required: (*i*) to evaluate the cell consumption rates of glucose, oxygen and lactate which are the three limiting substrates for energy production and (*ii*) to understand how these different consumption rates vary depending on the environmental conditions.

The production of ATP comes from two main processes and can thus be represented in two parts as proposed by Jagiella et al. [16]. The first part, is the ATP produced by glycolysis and the second one, the ATP produced by OXPHOS, if there is enough pyruvate and oxygen in the medium. Assuming that changes in glucose and oxygen concentrations in cells are primarily depending on their consumption and uptake rates, they write:

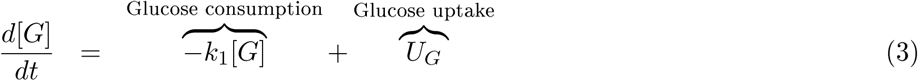

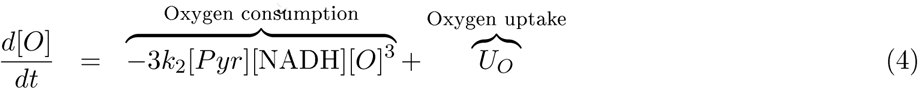

And, according to reactions (1) and (2), the evolution of the ATP concentration is given by:

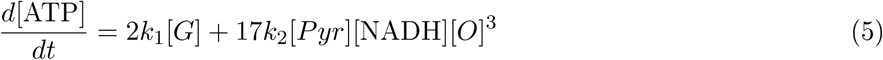

where [*G*], [*O*], [ATP], [*Pyr*] and [NADH] are the intracellular concentrations of glucose, oxygen, ATP, pyruvate and NADH respectively. *k*_1_ is the rate of glycolysis and *k*_2_ is the oxygen consumption rate through the citric acid cycle and OXPHOS combined. Considering the equilibrium condition where the uptakes of glucose and oxygen are equal to their respective consumptions, this equation can be rewritten as:

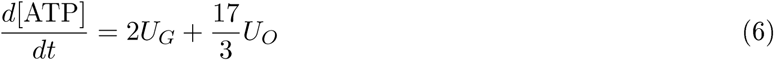

It is then required to evaluate the glucose and oxygen uptake rates *U*_*G*_ and *U*_*O*_ respectively. Casciari et al. [15] proposed a model, based on experiments on EMT6/Ro cells, where the uptakes depend on the concentrations of the substrates and on the pH. This model has been used and modified several times to describe the growth of spheroids [16] or to estimate the amount of ATP produced [11, 17]. However, the model does not integrate the energetic role of lactate. Only the acidification of the medium is taken into account. According to *observation 3*, this implies a significant drop in glucose consumption which raises the problem of the stability of the ATP level since it cannot be sustained long enough.

We constructed our model of cell metabolism by considering that the oxygen uptake is not directly depending on the glucose concentration by contrast with the other existing models [13, 15–17]. In the reference model of Casciari et al. [15], oxygen consumption (*via* OXPHOS) is directly linked to the level of glucose: the more glucose there is, the less oxygen is consumed (Fig 3A). In our model (Fig 3B), ATP is the factor that links oxygen to glucose: the more glucose is used to produce ATP, the less oxygen is consumed. This new hypothesis releases a strong constraint on the system and allows for more flexibility with the potential for generating more metabolic behaviors. *In vivo*, OXPHOS is not directly limited by ATP (however the reduction of ADP pool reduces its activity). But TCA enzymes like isocitrate dehydrogenase or oxoglutarate dehydrogenase are inhibited by ATP and NADH. By limiting these steps there is less NADH produced that can be used later for OXPHOS. Also, the less oxygen there is, the more glucose consumption increases. Indeed, when the cell lacks oxygen, HIF is stabilized and upregulates the expression of glycolytic enzymes [32–34]. This model improvement makes things more natural (i.e. more emergent).

**Figure 3:**
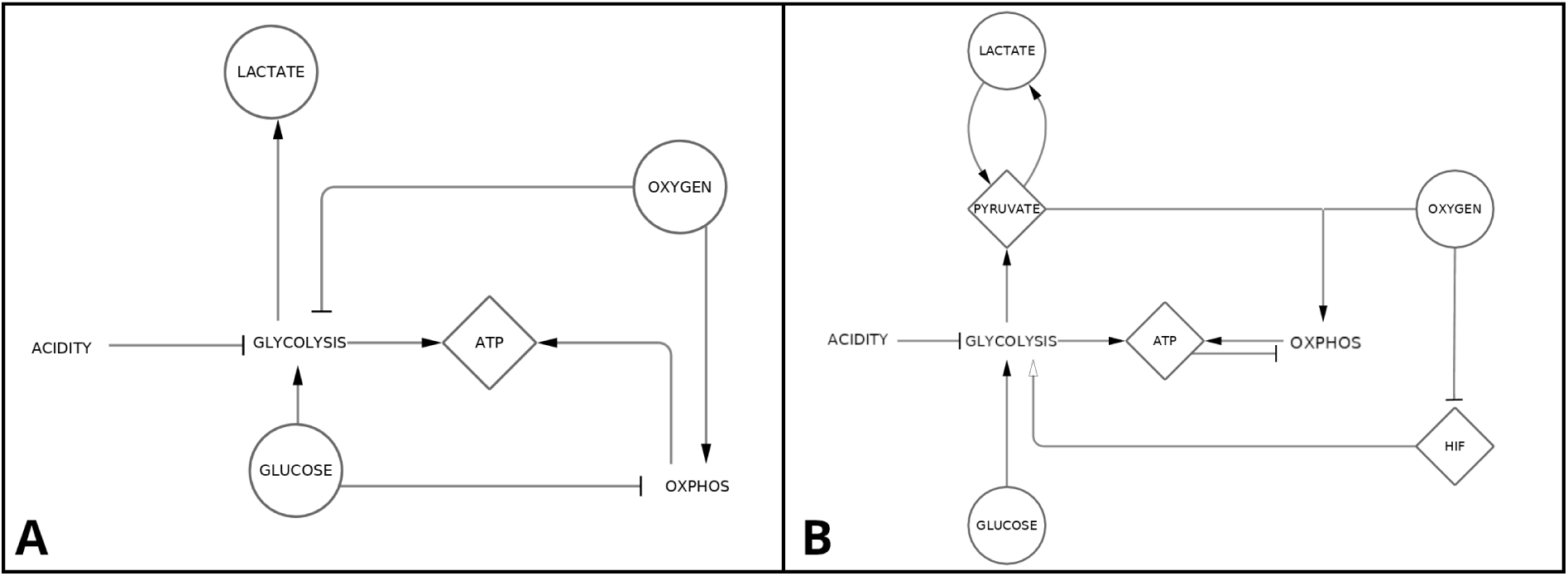
Comparison of the structure of the models built from Casciari’s model [15] with our model. **A.** Summary of models derived from the Casciari’s model. Some models only include certain elements, such as lactate [13, 16] or pH [17]. **B.** Overview of the model presented in this paper.

#### Glucose uptake rate, *U*_*G*_

According to the experimental *observation 1*, glucose uptake increases with extracellular glucose concentration up to a saturation threshold. The simplest way to represent this property is to use a Michaelis-Menten function:

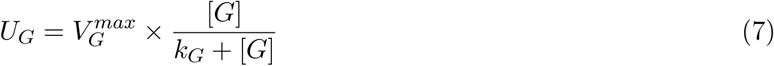

where *k*_*G*_ is the Michaelis constant for glucose consumption and 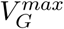 is the maximum uptake rate of glucose at saturation. First, the less oxygen there is, the more 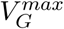 increases (*observation 2*). Additionally, the more acidic the medium, the more 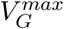 decreases (*observation 3*). This is also true when the pH becomes alkaline [35]. 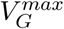 is thus expressed by the combination of these two effects as follows:

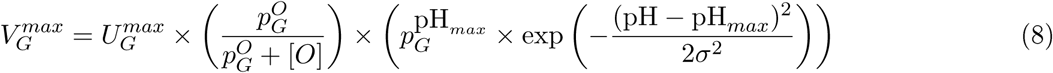

where 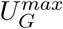 is the physiological uptake limit of glucose and 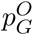 is a constant for glucose uptake variations according to the oxygen level. The pH-related term has a Gaussian form, which (*i*) varies from close to 0, when the pH is acidic, to 1 when it is physiological (pH ≈ 7.3), (*ii*) reaches a maximum at pH_*max*_, which is the pH corresponding to the maximum glucose uptake and (*iii*) decreases after. 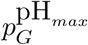 is the maximum expression of glucose uptake when the pH is optimum and *σ* is a constant that tunes the spread in the Gaussian term of the glucose uptake.

#### Oxygen uptake rate, *U*_*O*_

As for glucose and according to *observation 1*, the oxygen uptake is described with a Michaelis-Menten function:

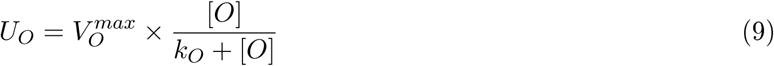

where *k*_*O*_ is the Michaelis constant for oxygen consumption and 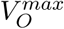 is the maximum uptake rate of oxygen at saturation.

Unlike glucose, the diffusion of oxygen across the plasma membrane is a passive diffusion. Thus, it can be considered that the concentration of oxygen at equilibrium, outside and inside the cell is almost identical. Only the oxygen consumption by the cell governs the inflow. The main role of OXPHOS is to provide the ATP needed for the cell and, the rate of ATP synthesis by OXPHOS is tightly coupled to the rate of ATP utilization [36]. Rather than varying the uptake of oxygen as a function of glucose concentration as in previous models [11, 15, 16], we hypothesize that it directly varies according to the need for ATP not filled by glycolysis. Indeed, there is no molecular evidence to link directly the evolutions of the two substrates. There are, in the other hand, a multitude of other signals that indirectly relate the two. If the cell needs a specific ATP level (ATP_*target*_), the mitochondria needs to produce the missing part of ATP (ATP_*target*_−ATP_*glycolysis*_) to complement the part produced by glycolysis (ATP_*glycolysis*_).

From (eq.2) and (eq.6), to produce 1 mole of ATP, the mitochondria needs 3/17 mole of oxygen. For 3 moles of oxygen one mole of pyruvate is consumed. Taking into account the fact that the level of pyruvate may be limiting, 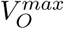 is expressed with the following expression:

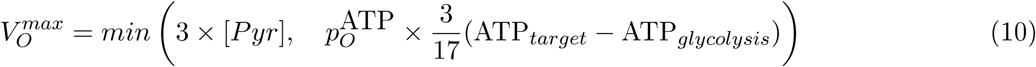

with 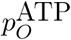, a sensitivity parameter for missing ATP.

#### Lactate uptake rate, *U*_*L*_

Lactate can be produced and secreted as well as consumed. Again, as glucose and oxygen, lactate uptake can be written with the following expression (*observation 1*):

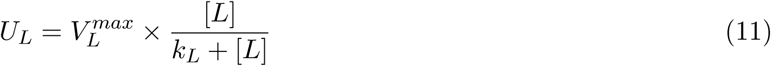

where [*L*] is the lactate concentration and *k*_*L*_ is the Michaelis constant for lactate consumption. Xie et al. [18], measured the amount of lactate consumed according to the level of lactic acidosis. They have shown that in low pH medium, the higher the extra-cellular level of lactate, the more the lactate uptake increases to the point where the inflow exceeds the outflow (*observation 4*). Lactate transport is done by monocarboxylate transporters (MCT), a group of proton-linked plasma membrane transporters. A proton gradient between the outside and the inside of the cell is required to transport lactate [37]. The parameter 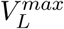 is therefore taken as a Hill function that decreases with increasing pH.

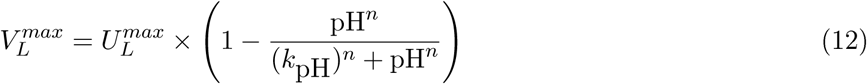

#### Pyruvate fate and Lactate secretion

The change in intracellular concentration of pyruvate is written as:

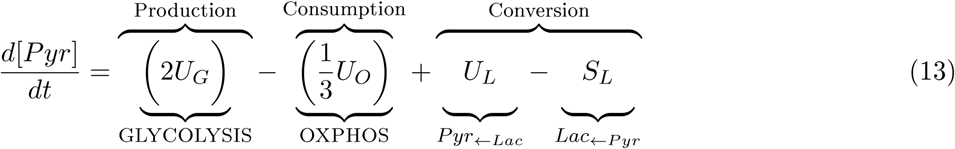

Xie et al. [18], observed that the level of pyruvate remains constant regardless of the pH and lactate conditions (*observation 4*). In this case the previous formula can be written as:

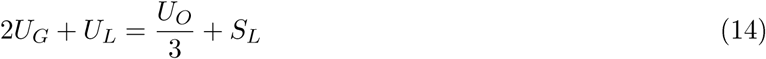

Finally, pyruvate converted to lactate corresponds to the “surplus” pyruvate, [*Pyr*]_*Target*_ being the basal concentration in the cell. Since this is a surplus, this term should not be negative:

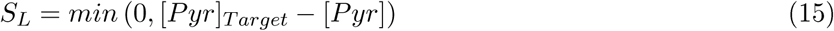

The functions defined in (eq.8–12) are fitted and parameterized from experimental data. Table 1 recapitulates the values used in the model and their sources.

**Table 1:** Parameters used to performed the simulations

| Symbol | Value | Description | References |
| --- | --- | --- | --- |
| $[G]$ | $[0, 15]$ mM | Extra-cellular concentration of glucose | [15] |
| $[O]$ | $[0, 0.1]$ mM | Extra-cellular concentration of oxygen | [15] |
| $[L]$ | 0 mM | Extra-cellular concentration of lactate | [18] |
| $[Pyr]$ | 0.12 mM | Intra-cellular concentration of pyruvate | [18] |
| $ATP_{target}$ | 2.8 mM | Intra-cellular concentration of ATP | [3] |
| $U_G^{max}$ | $1 \times 10^{-7}$ mol/cell/s | Physiological uptake limit of glucose. | Estimated<sup>1</sup> |
| $U_L^{max}$ | $1 \times 10^{-6}$ mol/cell/s | Physiological uptake limit of lactate. | Estimated |
| $k_G$ | 0.04 mM | Michaelis-Menten constant for glucose consumption | [15] |
| $k_O$ | $4.6 \times 10^{-3}$ mM | Michaelis-Menten constant for oxygen consumption | [15] |
| $k_L$ | 21.78 mM | Michaelis-Menten constant for reverse LDH reaction | [65] |
| $k_{pH}$ | 6.9 | pH at which the uptake of lactate is at half its maximum capacity. | Fitted from [18] |
| $p_G^O$ | 0.24 | Constant of glucose uptake variation according to oxygen level | Fitted from [15] |
| $p_O^{ATP}$ | 1 | Constant that defines the sensitivity to missing ATP | Estimated |
| $p_G^{pH_{max}}$ | 1.3 | Maximum expression of glucose uptake when the pH is optimal. | Fitted from [18] and [35] |
| $pH_{max}$ | 7.5 | pH at which glucose uptake is maximal. | Fitted from [18] and [35] |
| $\sigma$ | 0.27 | Constant for spread in gaussian term of the glucose uptake | Fitted from [18] and [35] |
| n | 67.95 | Hill coefficient for uptake of lactate according to pH. | Fitted from [18] |
| $ATP_{demand}$ | $[0.1; 0.2495; 1] \times 10^{-5}$ mol/cell/s | ATP needs for a cell from low to high removed from the pool at each time step | Estimated<sup>2</sup> |

<sup>1</sup>This parameter is set high to accelerate the simulation (this does not change the conclusions of the simulation) the value usually observed is around: $[0.1-1] \times 10^{-16}$ mM.s<sup>-1</sup>, see [15, 16].
<sup>2</sup>This parameter is also set high to match the high uptakes, the value usually observed is around: $[0.1-1] \times 10^{-15}$ mM.s<sup>-1</sup>, see [16, 17]

### Metabolic landscape depending on substrates availability

The model satisfies experimental observations. To test the model the extracellular concentrations were taken between 0 and 0.1 mM for oxygen and between 0 and 5 mM for glucose. Those values are compatible with *in vivo* concentrations. Uptakes have been normalized to facilitate the comparison between conditions. As oxygen becomes scarce or the extracellular glucose concentration increases, glucose uptake increases (Fig. 4A). Conversely, the presence of glucose lowers the uptake of oxygen (Fig. 4C). The effect of pH makes the uptake of glucose close to zero under acidic conditions (Fig. 4B). As a result, the uptake of oxygen no longer depends on the presence or absence of glucose in the medium at acidic pH (Fig. 4D).

**Figure 4:**
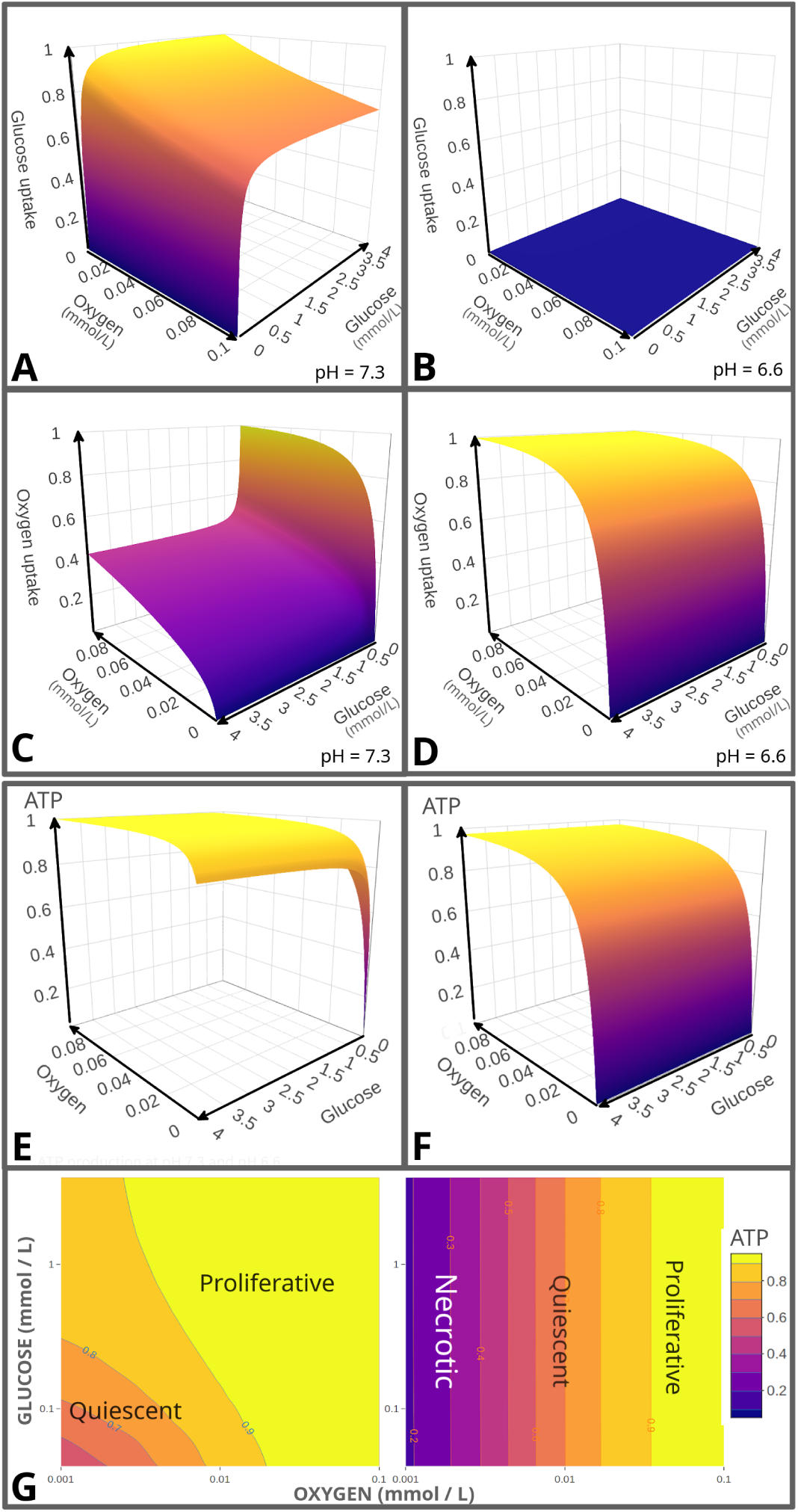
Normalized uptakes at different pH and extracellular concentrations of oxygen and glucose in mM: **A.** Glucose uptake at pH 7.3; **B.** Glucose uptake at pH 6.6; **C.** Oxygen uptake at pH 7.3; **D.** Oxygen uptake at pH 6.6. Normalized ATP production rate in multiple conditions: **E.** ATP Production rate at pH 7.3; **F.** ATP Production rate at pH 6.6; **G.** Heatmap of ATP production rate at pH 7.3 (left) and 6.6 (right) with a logarithmic scale. For each ATP level a cellular state can be associated. Typically low levels of ATP correspond to quiescent cells (reduced metabolism) whereas high levels of ATP are associated to proliferating cells.

From the previous uptakes, the production rate of ATP can be calculated (see eq.6). Several models do not take into account the need for ATP as a mechanism to regulate the uptakes of the main substrates. However, it is expensive for the cell to overproduce ATP [38] as it is disadvantageous to under-produce ATP when the surrounding resources are available. The result is that the cell cannot ensure a constant level of ATP as soon as the concentration of one of the extracellular substrates is changed. The cell needs a precise level in ATP and adapts its energy mechanisms to meet this need. Here, this is the OXPHOS that meets the ATP needs unfilled through glycolysis. Figure 4E shows that the cell − when the conditions are not extreme (concentrations of glucose or oxygen not close to zero) − can reach a plateau and stabilize its production of ATP to meet a fixed need. When substrates are close to zero, ATP production declines rapidly. In acidic conditions (Fig. 4F), the cell loses its dependence on glucose. The production of ATP depends mainly on the concentration of oxygen (and pyruvate not shown on this figure).

Figure 4G is a two-dimensional (2D) projection of figures 4E and 4F, with a logarithmic scale. This 2D representation is interesting to observe the levels of ATP production. Although it is not possible to reduce the cell state exclusively to its ability to maintain a certain level of ATP, it is clear that a proliferative cell uses more ATP than a quiescent one [39]. It is then possible to associate different cell types to different regions since the probability to encounter proliferative cells is bigger in high ATP production regions and necrotic cells in low ATP production ones.

### On the importance of the environmental conditions and cell energetic needs: no switch, nor reprogramming

The cell energetic metabolism is often reduced to an observation at a given time point, however it is a highly dynamical process. The cell constantly adapts to the changing environment and to its energetic needs (mass production, division, *etc*). It therefore makes sense to consider the temporal evolution of the system given the environmental context. We here consider three typical situations:

1. a non-limiting oxygen case compatible with 2D *in vitro* cell cultures;
2. a limiting oxygen case compatible with a poorly perfused environment;
3. a varying oxygen case that mimics tumor angiogenesis.

For each of these three different environmental conditions, we consider three different levels of cell energetic demand: low, medium and high.

Figure 5.A presents the *non-limiting oxygen case*, where oxygen is maintained at a constant level throughout the simulation. Glucose is, in the other hand, consumed with time and decreases with an intensity related to the cell energetic demand. We note that for the higher energetic demands, the glucose uptake saturates. This is why the drop in glucose concentration is the same for the medium and high ATP demands (both are beyond the saturation level). For the low energetic demand, OXPHOS is low, since there is no need for more ATP. This low OXPHOS is insufficient to absorb all the pyruvate produced through glycolysis. As a consequence, the excess pyruvate is transformed into lactate, which is excreted thus increasing the acidity. This pH drop creates a negative feedback on the glucose uptake until an equilibrium is reached between the glycolytic flux and OXPHOS. In other words the pyruvate production (through glycolysis) and consumption (through OXPHOS) become equal thus stabilizing the pH. This corresponds to a glycolytic contribution to ATP production of 5.5% (details are available in the Supplemental Information).

**Figure 5:**
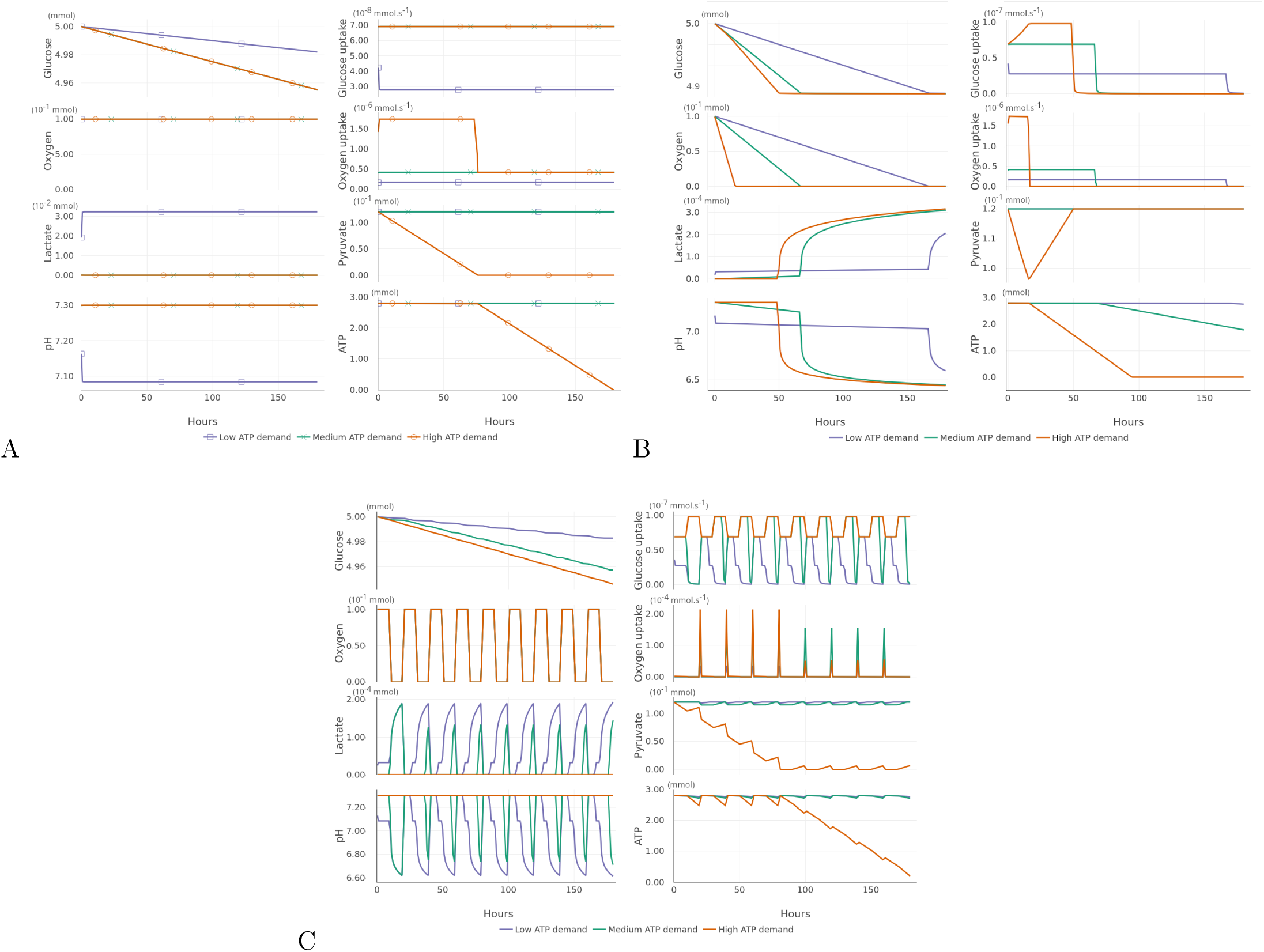
Evolution of the main metabolic molecules and uptake rates of glucose and oxygen as a function of the cell energetic demand from low to high. **A.** Oxygen is non-limiting; **B.** Oxygen is limiting; **C.** Oxygen is varying.

When the ATP demand is high, glycolysis is not sufficient to produce the pyruvate that feeds the OXPHOS flux for which the oxygen uptake is high. As a consequence, the pyruvate drops to extinction. Since OXPHOS cannot be fueled, the high level of ATP cannot be maintained and drops too. We note that this situation is rarely observed *in vivo* since the cell tends to reduce its energetic demand to avoid this situation. Moreover there are numerous alternative energetic substrates and pathways that the cell can use to produce its energy and that we have not considered here.

Figure 5.B presents the *limiting oxygen case*, where oxygen is rapidly consumed over time. There are three oxygen drop intensities that relate to the three levels of ATP demand. For the low ATP demand, we initially observe the same dynamic as in the previous case: the pyruvate production (through glycolysis) is higher than the pyruvate consumption (through OXPHOS). The surplus of pyruvate is converted into lactate and comes out of the cell with a proton until the pH is stabilized signing the equilibrium between OXPHOS and glycolysis. As soon as anoxia is reached, OXPHOS stops and pyruvate is entirely converted into lactate. Lactate is excreted and this leads to a second acidic drop.

For the high ATP demand, we note a sharp drop of pyruvate since OXPHOS consumes more pyruvate than glycolysis can produce. As soon as oxygen disappears, OXPHOS stops and the pyruvate pool is refilled. During the initial oxygen decrease, the glucose uptake increases transiently until saturation (limited uptake capability). This dramatically increases the acidity that ultimately results in the glucose uptake collapse.

Figure 5.C presents a case with *varying oxygen conditions* that can be considered as a proxy for tumor angiogenesis. It is well known that the angiogenic network is highly unstable leading to oscillating oxygen conditions with a wide range of periodicities from the second to several hours [40]. We chose here to simulate oxygen cycle with a periodicity of 20 hours. This periodicity has been chosen so as to enhance the observed effects and increase the contrast with the previous case.

For the high ATP demand, we again observe the same dynamic as in the previous cases. The pyruvate is entirely depleted by OXPHOS. At first the amount of pyruvate remains sufficient in the cell, despite its decrease, until it drops down. Small amounts are cyclically restored through glycolysis and are instantly consumed as soon as the level of oxygen allows it.

When the energetic load is lower, we observe sustained oscillations of lactate and pH caused by the periodic shutdown of OXPHOS. The oxygen oscillations frequency is fast enough to maintain an almost constant level of pyruvate restored by lactate. We note that we did not take into account the time needed to convert lactate into pyruvate − in our model it is considered as an instantaneous process − this leads to sharp pH oscillations that are not experimentally observed. However our aim at this stage is to highlight the possible emerging metabolic behaviors rather than being quantitatively realistic.

These results clearly show that − depending on the oxygen constraint − glycolysis and OXPHOS cooperate to sustain, as far as possible, the energetic demand in terms of ATP production. This cooperation is mediated by the amount of pyruvate which is the product of the first and source of the second. This contradicts the *switch mechanism* of the metabolism that implies an alternating and exclusive (*i.e.* dual) functioning of the two metabolic modes. Our results also show that the cell does not need any *metabolic reprogramming* to adapt to the changing environmental constraints − at least on the short term − to maintain its energetic needs, according to the definition of this process that can be found in the literature [41–45]. These two notions will be thoroughly discussed, at the light of our results, in the dedicated section.

### Why cell metabolism measurements at the cell population scale can be misleading

The spatial dimension is often neglected in most studies on cell metabolism including theoretical approaches and experimental ones. In the first case, most models focus on the mechanisms on the individual cells (as we did it until now) and in the second experimental case, most measurements are realized at the level of the entire cell populations either on two dimensional cell cultures or three dimensional spheroids. Extracellular Flux Analysers (Seahorse) [46] are widely used to characterize cell metabolisms based on the measures of OCR (Oxygen Consumption Rate) and ECAR (Extracellular Acidification Rate). However these measurements reflect mean values for the entire cell population, when in fact high discrepancy exists between cells depending on the local environmental context and cell state [47, 48]. If the use of this device makes more sense for two dimensional cell cultures where the environment is supposed to be homogeneous, it is clearly not adapted for spheroids where the inner cells do not have the same access to the resources compared to the peripheral cells.

In this section we aim to specifically highlight the heterogeneity that exists in a spheroid due to the gradients of the main substrate from the periphery to the core (Fig. 6, upper graph). These gradients are essentially due to the increased density of cells in the centre that consume more nutrients and impede their diffusion. As a consequence, the gradients of resources (oxygen and glucose) induce the emergence of different metabolic states depending on the cell depth in the spheroid. We performed an illustrative simulation presented in figure 6. The initial state corresponds to a tumor spheroid immersed in a highly hypoxic environment. To highlight the different metabolic states that can be encountered we have not considered cell states transitions from proliferation to necrosis. However we take into account the increase cell cycle duration induced by the lack of oxygen mediated by HiF-1*α* [49]. Details on the model are given in the Supplementary Information.

**Figure 6:**
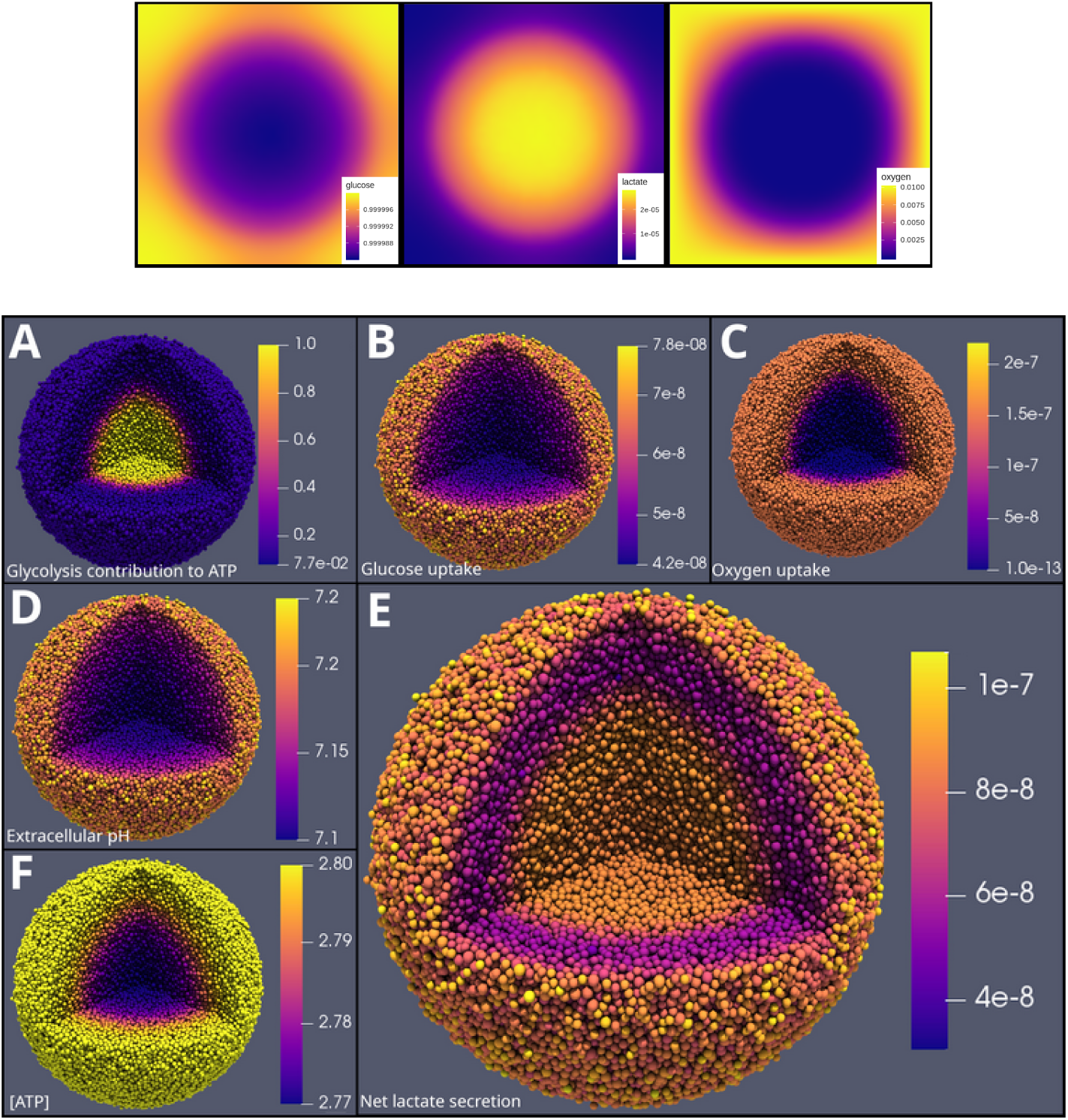
Upper graph: distribution of molecules and acidity in the medium of the spheroid simulation. From left to right: Glucose concentration in mM (initial concentration in the medium: 1mM); Lactate concentration in mM (initial concentration in the medium: 0mM); Oxygen concentration in mM. The oxygen concentration is fixed (0.01 mM) at the boundary of the simulation domain. Spheroid simulation at 10 days using Physicell[50]. Initial medium concentrations: 2mM glucose, 0.01mM oxygen, pH=7.3, no lactate. Each figure represent the same spheroid with coloration for different parameters. **A.** Glycolysis contribution to ATP production; **B.** Glucose uptake in *mM.s*^−1^; **C.** Oxygen uptake in *mM.s*^−1^; **D.** Extracellular pH; **E.** Net lactate secretion in *mM.s*^−1^; **F.** ATP level in mM. Movies of the simulation are available in the Supplemental Information.

Figure 6A clearly shows that the glycolysis contribution is much higher in the centre where the oxygen level is low (Fig. 6, upper graph). As a consequence, the pH is lower in the centre and is directly correlated to the glucose uptake gradient (Fig. 6D and 6B). In the other hand, the gradient of oxygen uptake is steeper from the centre to the periphery since the consumption of oxygen is higher than glucose and its initial concentration is lower making it more sensitive to depletion (Fig. 6C). The net lactate secretion exhibits an interesting profile (Fig. 6E) with a mid layer of lower secretion. This is explained by a high OXPHOS activity requiring pyruvate at the outer layer. This leads to a lower level of secreted lactate since glycolysis is slightly diminished from the periphery. At the other end, in the centre, OXPHOS is highly diminished since oxygen level is low. This reduces the need for pyruvate (by OXPHOS), therefore excess pyruvate is converted into lactate. The net lactate secretion thus reaches almost the same level at the heart of the spheroid than at its periphery. Finally, ATP is globally maintained around its basal level, except at the centre where the oxygen is low and OXPHOS is reduced (Fig. 6F). We note that glycolysis is not able to sustain the ATP level. This would typically induce a transition towards a reduced metabolism such as quiescence. However, the cell usually reduces its energetic needs much before it lacks ATP, since HIF triggers the entry into quiescence as soon as the oxygen level is too low.

Concerning the Warburg effect, according to the classical definition, it corresponds to the production of lactate in presence of oxygen. Figure 6E, that shows the net lactate production, exhibits a non-homogeneous Warburg effect. Its intensity − defined by the importance of the glycolysis contribution (Fig. 6A) − depends on the gradient stiffness of the substrates. Moreover we here observe one instant of an evolving process that corresponds to a transitory state. This shows that this effect is not a well-defined state with a switch-like dynamics but a gradual event. We note that our simulation is only one possible configuration given our choice of parameters. Other cases could be generated. For example a sharper pH gradient from centre to periphery leading to extinction of glycolysis would have led to different cell metabolism scenarios.

## Discussion

The cell metabolism is highly complex since it is a multifactorial mechanisms that involves many different interacting processes with many different actors. Moreover it is an evolving process and although crucial, this aspect is rarely considered and often overlooked.

In this context, a theoretical approach is well adapted since it allows to make sense of this complexity and to address the temporality. It allows to test the pertinence of some new hypotheses so as to provide a better understanding of the intimate functioning of the metabolic machinery and to provide new insights to guide future research.

The model of cell energy metabolism that we proposed in this study integrates the most recent knowledge. It is based on a number of key experimental observations and established facts. It is moreover fitted and parameterized from experimental data.

The model focuses on the glycolysis-OXPHOS relationship and emphasizes in particular the role of lactate as a substrate, as well as the central role of pyruvate in the regulation of the metabolism. The latter makes the link between glycolysis, fermentation and OXPHOS (after conversion in the TCA cycle).

The oxidation of pyruvate requires that it is imported into the mitochondrial matrix and subjected to the activity of the complex pyruvate dehydrogenase (PDH). The activity of this enzyme is regulated by several conditions, such as CoA levels, NAD+/NADH ratio. It is a relatively long process in comparison with fermentation. On this particular point, we note that glycolysis is very often depicted as *less efficient* than OXPHOS [1, 8, 51, 52]. This statement is based on the amount of glucose consumed per molecule of ATP produced. By relying exclusively on this observation it seems obvious that glycolysis is less efficient than OXPHOS. However, in terms of ATP molecules produced by unit of time, glycolysis is a much faster process (about a hundred times faster), and hence the most efficient to produce energy. It allows the cell to adapt quickly in order to meet immediate energy needs. Glycolysis is not efficient with respect to the amount of glucose consumed but it is more efficient than OXPHOS to produce some energy very rapidely to respond to acute needs [28, 53].

When the glycolytic flux exceeds the maximum activity of PDH, the cell spontaneously transforms the pyruvate into lactate in order to get rid of the excess amount. [51].

It is also interesting to note that while glucose consumption and lactate production decrease with pH, the concentration of intracellular lactate increases. This mechanism suggests that the cell recovers extra-cellular lactate to maintain its level of pyruvate at a constant value. Indeed, in lactic acidosis conditions, the protons level being higher outside the cell, the entry of lactate via the transporter MCT1, is facilitated [18].

We could picture this system as a hydraulic dam. The dam retains upstream pyruvate, which founds its source in glycolysis. Depending on the energy needs, the dam is open with more or less intensity (OXPHOS). Sometimes the level overflows, the dam then lets pass the surplus without producing energy. Pyruvate is then transformed into lactate. Conversely, when pyruvate is lacking, the dam is supplied through the lactate source.

The simulations we performed to observe how imposed environmental constraints (*i.e*. the oxygen level and acidity) and imposed energy requirements push the cell to adapt, highlight this dam mechanism. Our results clearly show that glycolysis and OXPHOS are used concomitantly and in a cooperative way [20], with a gradation in their relative contributions to ATP production that depends on the available resources and pH.

These results somehow contrast with the current vision of the tumor cell metabolism, which is depicted as abnormal and characterized by an increased glycolysis even in presence of oxygen, the so-called Warburg effect. It seems important to us to stick to the original definition of the Warburg effect, that does not presuppose the presence of oxygen [5, 6, 53]. This effect − *i.e*. the suproduction of lactate − is acknowledged as a hallmark characteristic of cancer. However, recent experiments have shown that by acidifying its environment the tumor cell progressively auto-inhibits this phenotype [18]. By integrating this observation into our model, our simulations show that the Warburg effect is indeed only transitory (time-limited) and contextual (acidity-dependent). Therefore it is not an inherent characteristic of the tumor cell, but a spontaneous and transitory adaptation mechanism not fundamentally different from a normal cell. In response to Upadhyay et al. [54], this effect appears more as an epiphenomenon that plays no causal role in tumorigenesis.

Indeed, the increased glycolytic activity in the presence of oxygen is not a phenomenon specific to cancer. There are brain regions that primarily use aerobic glycolysis [55], as well as endothelial cells during angiogenesis [56], and mesenchymal stem cells that rely primarily on glycolysis and require less oxygen [57]. It might be useful to recall that, from the evolutionary point of view, OXPHOS is a process that appeared after glycolysis. The primitive eukaryotic cells became able to use oxygen only after the endosymbiosis of proteobacteria. The eukaryotic cells have thus obtained by this mean an additional source of ATP production, complementary to glycolysis, and that has been preserved until now.

In this context, the expression *metabolic switch*, often used to express the tumor cell metabolic change, appears inappropriate. This expression is indeed emphasizing a dual nature of tumor metabolism, thus reinforcing the idea that the cell would either use fermentation or mitochondrial respiration as two opposite, non-concomitant mechanisms, which is a distortion of reality.

In a similar way the notion of *metabolic reprogramming* − often used in the recent literature [42–45] − can also be misleading. According to some definitions, *metabolic reprogramming* can refer to “*the ability of cancer cells to alter their metabolism in order to support the increased energy demand due to continuous growth, rapid proliferation, and other characteristics of neoplastic cells*" [41]. However, this definition is too vague and appears inadequate since our model responds to it, by allowing the cell to adapt, without reprogramming. Moreover, it is important to note that there is no alterations as such in the tumor cell metabolism: the metabolic pathways are modulated − overexpressed or underexpressed − but they are still fully functional. Until now, no new alternative metabolic pathways have been identified that would justify to treat the tumor metabolism as structurally abnormal.

Therefore, *reprogramming* necessarily implies that the origin of the tumor metabolic adaptations, is either genetic (acquired mutations) or epigenetic (genes expression) for the cell to acquire its enhanced metabolism [58]. Many proto-oncogenes or tumor suppressor genes are involved in the expression of signal transduction pathways and may have an impact on metabolism, for example by increasing the expression of glucose transporters on the cell surface via PI3K/Akt pathway [42]. In our model this would lead to an increase in the glycolytic capacity associated with the glucose uptake. While these factors may modulate the activity of metabolic pathways, they do not change their basic mechanisms, an idea also mentioned by Bhat et al. [2] as *metabolic plasticity*.

Temporality, and more specifically the time scale with which changes in the metabolism develop, has been overlooked despite its importance. On a short time scale, our results contribute to show that the early tumor cell is able to gradually adapts its metabolism and the metabolic changes are reversible [28]. On a longer time scale, metabolic reprogramming occurs and it most probably depends on the stress history (frequency and intensity of hypoxic events) that the cell has experienced. It is well known that HIF*−*1*α* stabilization with hypoxia is a powerfull driver of genetic instability [59]. To our knowledge no studies have been realized yet to trace and characterize the evolution of the cell metabolism − at the individual cell scale − during the course of tumor development. Such studies would help to decipher the level of environmental constraints required to induce reprogramming.

At this stage our model focused on the short time scale metabolic adaptation. The model was developed to manage the complexity of the different reactions and to make sense of these reactions in space and time. Our results first emphasize the cooperativity of the metabolic modes glycolysis and OXPHOS. This disqualifies the notion of a *metabolic switch* by showing that the Warburg effect is not an inherent characteristic of the tumor cell, but a spontaneous adaptation mechanism not fundamentally different from that of a normal cell. In a second time, we clarified the related notion of *metabolic reprogramming*. We argued that the tumor metabolic pathways − although modulated − are fully functional. We therefore reached the conclusion that the tumor cell energy metabolism might not be that abnormal, within the limit of the glycolysis-OXPHOS relationship that we explored.

By extrapolation to the other metabolic modes, if there is indeed no structural metabolic defaults in a tumor cell that would allow to differentiate it from a normal cell − *i.e*. only modulated metabolic pathways, but no new nor destroyed pathways − then we can ask: is it really pertinent to look only for a therapy targeting [60] mutated metabolic genes in tumor cells ? Considering that there is no fundamental differences between normal and tumor cell metabolisms suggests that normalizing the environment could be a good strategy [61, 62]. Since the tumor cell metabolism is identified as abnormal due to the disregulated environmental context, the return to normality (in terms of oxygenation and pH) would stabilize the cells rather than enhance their genetic instability and consecutive aggressiveness.

More refined studies, specifically focused on the the evolutive aspects of cell metabolism during tumor development, would be useful to address this issue and explore such alternative therapeutic strategies.

### Limitation of this study

The main difficulty encountered when building a theoretical model is its parameterization. Not all parameters come from the same experiments. They are obtained for different cell lines and with different protocols which implies some variability on each parameters. The choice of the parameters used in this study is thus based on a consensus. As a consequence, our aim was limited to observe the qualitative emergence of different metabolic behaviors depending on some imposed conditions related to the environment and cell energetic needs. To reach a quantitative accuracy, it would be necessary to acquire more fundamental data on the consumption rates of the cells for the different substrates and to establish energetic profiles in a standardized way, at the level of several cell lines, as indicated by Zu and Guppy [53].

We are aware of the tremendous complexity of the cell metabolic machinery. However we deliberately chose to simplify it, by focusing on the interactions of three main metabolites (glucose, oxygen, lactate) allowing to reduce the computational cost with the idea to look at their impact at the tissue scale. As a consequence, we proposed a relatively simple model − for which we already explained some of the limitations above − and we have additionally ruled out the following mechanisms:

1. most of the responses related to cellular or environmental changes are instantaneous, for example the inter-conversion between pyruvate and lactate. This would not make any fundamental differences in our results, but this would slightly delay some emerging behaviors.
2. The mitochondrial responses to ATP needs is limited only by the availability of the substrates and not by the mitochondrial mass. The model is therefore limited to the short time scale cell response and does not integrate longer time scale adaptations involving mitochondrial biogenesis.
3. The model has been focused on the two main metabolic pathways: glycolysis and TCA-OXPHOS to form a core model. The other metabolic pathways − *β*-oxydation, glutaminolysis, ketogenesis − have not been considered at this stage. However they can be added to the core model.
4. There are multiple sources of acidity and we only considered the acidity from the transport of lactate. While it helps to focus on the specific effect of lactic acidosis it may neglect other important acidity sources like CO_2_
5. The balance between biosynthesis of amino-acid and energy production has not been considered.

The consequence of these limitations on the results, is that we highlight major behaviors and qualitative tendencies. More precise models would not dramatically change the results but would allow to reach a quantitative description and test fine regulation mechanisms.

## Methods

### Cell energy metabolism simulations

To simulate the evolution of metabolic molecules (Fig. 6-8), we used the *DifferentialEquations.jl* module (v 1.10.1) [63] of the Julia language (v 1.1.1). This module integrates many Ordinary Differential Equations (ODE) solvers and we have chosen the solver *AutoTsit5-Rosenbrock23*, which allows to automatically choose the algorithm adapted to the stiffness of the problem (more details are available on the documentation page of the module): https://docs.juliadiffeq.org/latest/solvers/ode_solve.html.

The julia code is available in jupyter notebook format (.ipynb) at the following link: https://mycore.core-cloud.net/index.php/s/fJnOfsNPgYjpmOh

### Spheroid simulation

To produce the spheroid simulation, we have integrated our model of cell energy metabolism as a module in PhysiCell (v1.5.2 available at http://physicell.mathcancer.org/) [50], an open source physics-based cell simulator which manages, among other things, the diffusion of substrates in the culture medium and gives tools to define the cell cycle, division rates, necrotic and apoptotic events. As an agent-based model, each cell is independent and has its own internal processes (cell cycle, energetic metabolism). They interact with each other and share local resources. In our model implementation, the diffusive resources are: oxygen, glucose, lactate and protons. We have integrated each phase of the cell cycle. The standard cell cycle duration is fixed at 24 hours (G1: 11h, S: 8h, G2: 4h, M: 1h) [64], in a medium with optimal oxygenation (fixed at *pO*_2_ = 38mmHg, or 0.05282 mM). If oxygen is missing the duration of the G1 phase extends proportionally to the ratio: *pO*_2_/38.

### Model parameters table

## Acknowledgements

This work is related to the *POET project* financed by INSERM/CNRS “Santé Numérique − 2019" (Grant Reference: 216669). We thank the Research Group GDR ImaBio (http://imabio-cnrs.fr) for partial funding of P.J.’s work. We also wish to warmly thank Paul Macklin for his help and support with PhysiCell that we used to make the spheroid simulation.

## Author Contributions

A.S. conducted this research and supervised the model development; P.J. developed the model, performed the simulations and produced the graphical representations. Both authors contributed equally to the writing of the paper.

## Supporting Information

### Contribution of glycolysis to the production of ATP not involving lactate

In order to keep the level of pyruvate constant, the pyruvate production must be equal to the pyruvate consumption (eq.14). To find the level of glycolytic ATP contribution that does not require any lactate, we use (eq.14) that gives:

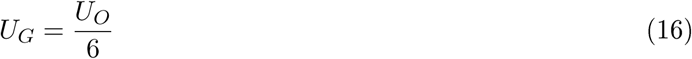

This relation is used in (eq.6) which becomes:

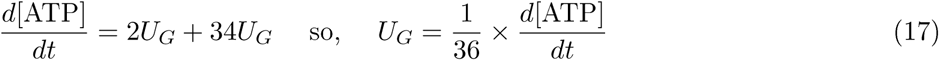

Since the ATP produced through glycolysis is obtained from:

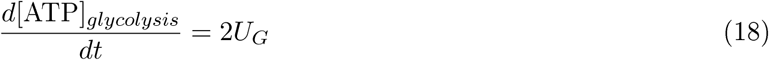

Then the combination of (eq.17) and (eq.18) gives:

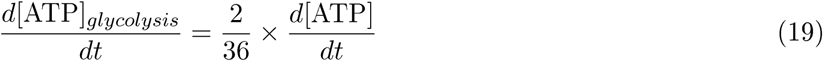

In conclusion, considering only glycolysis and OXPHOS as pathways involved in ATP production, if the glycolytic contribution to ATP is equal to 2/36 *i.e.* 5.55%, no lactate is produced and the level of pyruvate remains constant.

### Movie 1 - Spheroid simulation: evolution of oxygen uptake

3-day growth simulation of a mature spheroid using Physicell [50]. Initial diameter of the spheroid: 800*µm*. Initial medium concentrations: 2mM glucose, 0.01mM oxygen, pH=7.3, no lactate. The color scale corresponds to the oxygen uptake of each cell.

https://mycore.core-cloud.net/index.php/s/PbKUGggO9Sgoby4

### Movie 2 - Spheroid simulation: evolution of lactate secretion

6-day growth simulation of a mature spheroid using Physicell [50]. Initial diameter of the spheroid: 800*µm*. Initial medium concentrations: 2mM glucose, 0.01mM oxygen, pH=7.3, no lactate. The color scale corresponds to the net lactate secretion of each cell.

https://mycore.core-cloud.net/index.php/s/vDl4DBH79TKQpOe

### Figures

All figures are available at high resolution at the following link: https://mycore.core-cloud.net/index.php/s/1g9qtDlHYFBQRTb

